# Lipid-Driven Biophysical Selection of β-Furanoside-5’-Phosphate as the Sole Scaffold of Extant RNA

**DOI:** 10.64898/2026.05.26.727819

**Authors:** Ze-Run Zhao, Qian-Qian Chen, Hao-Xing Xu, Bing-Yue Zhao, Cheng-Cheng Gu, Xiao Wang

**Affiliations:** School of Chemistry and Chemical Engineering, Nanjing University, 163 Xianlin Avenue, Nanjing, 210023, Jiangsu, P. R. China

## Abstract

The emergence of RNA’s primordial backbone presents a fundamental question in the study of life’s origins: why was β-furanoside-5’-phosphate selected over other isomeric alternatives as the predominant building block during the origin of life? In contrast to previous studies that have demonstrated selective synthetic routes to canonical RNA structures, we explore biophysical factors that selectively favor β-furanoside and 5’-nucleotide over their undesired isomers. Inspired by the chromatographic elution order of nucleoside and nucleotide isomers, we establish a new selection model to discriminate between them. β-Furanoside is found to be more lipid-permeable than any other configurational isomer, making it the most abundant species after permeation. Outward permeation selectively screens intracellularly formed nucleotides, enriching 5’-nucleotide as the primary RNA building block within a protocell. The unique properties of representative canonical nucleosides and nucleotides are rationalized based on both their structural and dynamic characteristics, as elucidated by DFT and MD calculations. A scenario in which β-furanoside selectively accumulates in a lipid droplet is further investigated using a specifically designed micromixer. Together, these findings suggest the existence of a primitive selection mechanism driven by purely biophysical forces, which may have played a critical role in advancing key steps in the progression of primitive biomolecules.

## INTRODUCTION

The emergence of life and the required conditions to sustain it are believed to begin from Hadean or early Archean.^1^ The progression of primitive biomolecules is regarded as the foundation for complex life-related macromolecules and sophisticated metabolic systems. ^2^ RNA is hypothesized to be the first biological macromolecule that carries genetic code as well as catalyses early biochemical transformations. ^3^ The unique structure of canonical RNA came into being through a series of transformations and directional processes,^4^ or through evolution.^5^ A plausible abiotic synthesis of ribonucleotides is considered crucial for realizing the RNA World, yet several critical issues remain unresolved:^6^ (1) the selective formation of ribose or its skeleton; (2) the construction of *N*9-purine and *N*1-pyrimidine glycosidic linkages; (3) the selection of β-furanoside as the configuration of ribonucleosides; and (4) the regioselective phosphorylation. These problems pose persistent challenges to the RNA World hypothesis.

Traditionally recognized sugar-forming transformations (e.g. the formose reaction) afford ribose in low yield and selectivity.^7,8^ However, the necessity of free ribose can be circumvented with an elegant stepwise strategy developed by the Sutherland lab, in which pyrimidine nucleotides are selectively prepared with the desired β-*N*1-furanoside configuration. ^9^ Pyrimidine β-ribofuranoside can be exclusively prepared via the addition of crystalline *ribo*-aminooxazoline and cyanoacetylene, followed by thiolysis, photoanomerization and hydrolysis. ^10^ Canonical purine ribonucleosides can be selectively synthesized from thioanhydroadenosine via UV irradiation in the presence of sulfite.^11^ To seek a compelling evidence for the co-evolution of RNA and DNA, Sutherland later proposed the coexistence of purine deoxyribonucleosides and pyrimidine ribonucleosides as the precursors of primitive genetic polymers, in which the two classes of nucleosides share the same synthetic route.^12^ To address the limited availability of ribose, we developed a metal-doped-clay model, ^13,14^ with which ribose can be selectively enriched and stabilized from a complex, unselective formose mixture. For nucleosidation with free ribose, the Carell lab reported a landmark synthesis via the ribosylation of formamidopyrimidine and *N*-isoxazolylurea,^15,16^ which unambiguously produces *N*9-purine and *N*1-pyrimidine nucleosides. Borate was found to promote the accumulation of furanosides^15,16^ and the 5’-selectivity in the subsequent phosphorylation.^17,18^ The Trapp group developed a highly regio-and stereoselective path to prepare deoxyribonucleosides from canonical nucleobases, acetaldehyde and sugar precursors.^19^ Our lab demonstrated a cavitation-facilitated method to afford an elevated amount of *N*9-purine ribonucleosides.^20^ More recently, we discovered that titanium minerals catalyze the ribosylation of purines and pyrimidines with high regioselectivity for the *N*9 and *N*1 positions.^21^ However, a major challenge for most of these direct coupling strategies utilizing free ribose is that four isomers (α-furanoside/**α-f**, β-furanoside/**β-f**, α-pyranoside/**α-p**, β-pyranoside/**β-p**) are obtained, with the desired β-furanoside being a minor product because ribose mainly exists in β-pyranose form in aqueous media.^22^

Most of the aforementioned routes offer plausible solutions to Problems 1 and 2, but few attempted to address Problems 3 and 4. One key question remains: can the selection of the canonical β-ribofuranoside-5’-phosphate take place prior to its incorporation as an RNA monomer (Fig. 1)? First, are there mechanisms that selectively remove the undesired isomers formed alongside β-furanoside? Second, although several phosphorylation studies have reported elevated yields of 5’-nucleotide, ^23,24^ are there effective routes for removing the unwanted regioisomers (Fig. 1)?

**Fig. 1.**
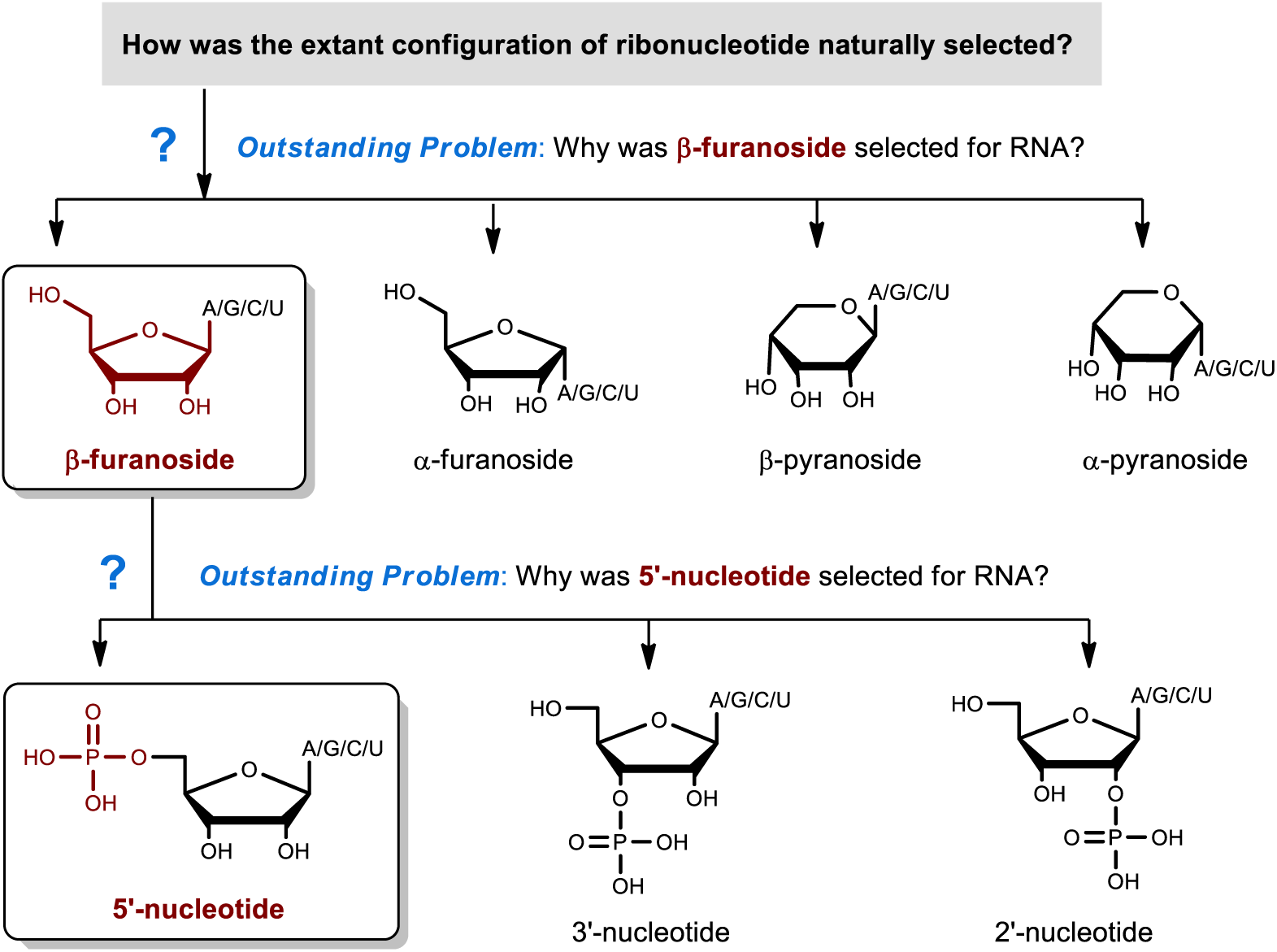
Challenges of the selection of the desired β-furanoside and 5’-nucleotide configurations (nucleobases: A, adenine; G, guanine; C, cytosine; U, uracil).

In response to this challenge, the ultimate establishment of the canonical ribonucleotide structure probably needs extra enhancement. In addition to the aforementioned routes, the configuration of RNA is believed by some biochemists to be the outcome of evolution, in which the extant RNA strand has conformational superiority due to its thermodynamic stability, as compared with genetic polymers with other configurations and linkages. ^25, 26^ However, in a polygenesis point of view, it is reasonable to predict the existence of other driving forces that promote the selection of monomer prior to oligomerization. In particular, physical processes such as transportation and separation^27,28^ deserve attention in understanding the selection of primitive molecules, as they might elevate the selectivity or purity of the target molecule by phasing out side products. Therefore, we set out to establish a selection model driven by such physical forces.

Even with a reasonable selection mechanism, product stability constitutes another worrying obstacle. ^29^ Nucleosides with different nucleobases have disparate levels of acid and base tolerance.^30^ Although the emergence of life may have taken hundreds of millions of years, the products of each synthetic step must have been stabilized and utilized in a timely manner.^31^ To avoid decaying over time, once formed, they should preferably seek stabilization on a solid support (e.g., a mineral surface),^32,33^ or enter a confined space with an unfluctuating environment (e.g., a protocell).^34,35,36^ Protocell has demonstrated efficacy in the selection of ribose,^37^ nonenzymatic template-directed RNA synthesis ^38^ and template copying. ^39^ However, to our knowledge, the potential role of protomembrane in selecting the canonical configuration of nucleoside and nucleotide has never been explored. There was one report on the permeation of a single nucleotide across the protomembrane, ^40^ but not on the comparison of permeabilities of different nucleoside/nucleotide isomers, nor about the selection of their extant configurations. Herein, we reveal that a lipid membrane or droplet possesses an intrinsic separation capability that promotes the selective enrichment of β-ribofuranoside and 5’-phosphate, the configuration required for RNA.

## RESULTS AND DISCUSSION

### Exploring the Selection of β-Furanoside with a Permeability Assay

When pondering over the question on whether any unique biophysical nature of β-furanoside might exist as the basis of it being separated from other isomers, we serendipitously noticed that for all genetic alphabets, β-furanoside is always the *longest* retained nucleoside isomer on a reverse phase (RP) HPLC column. This observation urged us to systematically look into the problem. The retention factors of configurational isomers of ribonucleosides were therefore measured (Fig. 2a; see Supplementary Information for details). The stationary phase of an ordinary RP-HPLC column comprises of silica modified with C_4_–C_18_ aliphatic chains. Therefore, we suspected that this distinctive elution pattern probably unveiled the uniquely high lipophilicity or membrane permeability of β-furanoside versus other configurations, which could be a plausible origin of its rise to prominence. Plausible protomembrane components include decanoic acid,^41^ amino acid stabilized-decanoic acid,^42^ oleate,^43^ polycyclic aromatic hydrocarbons (PAH)-incorporated fatty acid, ^44^ a combination of fatty acid/alcohol/glycerol monoester, ^45^ or phospholipid. ^46,47, 48^ Fatty acids and other oxygenated alkanes could be produced via the Fischer−Tropsch-type process under hydrothermal conditions.^49^ Among the less investigated non-fatty acid components, glycerol-1-phosphate is a plausible lipid precursor for the common origin of RNA, protein and lipid via a cyanosulfidic protometabolism.^50^ Regardless of lacking a solid answer of the exact composition of protomembrane (e.g., length of aliphatic chain, hydrophilic head group), it is reasonable to envisage the possibility of the selection of β-furanoside at a generic lipid barrier.

**Fig. 2.**
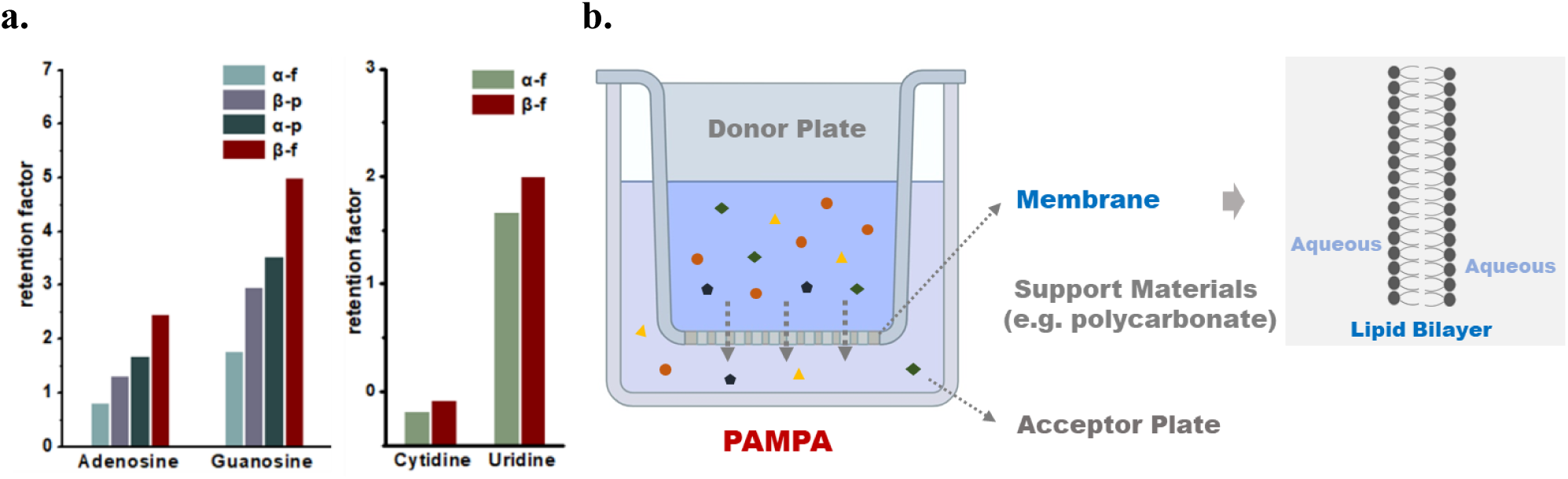
Foundation of a plausible selection pathway based on permeability, and the assay to explore the permeability order. (**a**) measured retention factors of ribonucleoside isomers (see Supplementary Information for details); (**b**) graphical illustration of PAMPA. Retention factors of nucleoside isomers were calculated by their retention times measured on a Shimadzu LC-2050 system with SymmetryShield^TM^ RP18 column (5 μm, 100 Å, 4.6 mm I.D. × 150 mm), at a detection wavelength of 254 nm, with water (adjusted with 0.1% formic acid) as the eluent at a flow rate of 1.0 mL/min.

Over the past two decades, several studies on the permeabilities of life-related molecules across protomembrane were reported.^37,40,45^ In most examples, permeability was measured by encapsulating one analyte and observing its leakage over time, utilizing size-exclusion chromatography to isolate the vesicles and fluorescence to monitor their volume.^51^ Our study faces a specific challenge: nucleoside isomers must be assessed in mixtures, while conventional protomembrane methods are designed to measure the permeability of a single compound at a time. Related studies raised a caution that the permeability of a compound may vary, depending on whether it is measured individually or with other compounds. For example, the transport of the anthelmintic drug praziquantel was found to be assisted by glycyrrhizin across a membrane made of 1,2-dioleoyl-sn-glycero-3-phosphocholine (DOPC).^52^ Besides that, there are also examples of competitive permeation. At slightly elevated temperatures in gas separation using polymeric membranes, the higher activity of H_2_O leads to an increase in its permeability and a significant decrease in those of CO_2_ and CH_4_.^53^ In addition, the reported vesicle permeability of a compound is usually calculated from a standard curve plotted with an impermeable reference solute, rather than with the compound itself. For example, the permeabilities of sugars and alditols were determined by monitoring the volume change in vesicles after adding the compound to the extracellular fluid, in which a spectrofluorometer was used, with fluorescence data converted into volumes based on a standard curve obtained from osmotically compressed vesicle experiments. ^37^ The permeability coefficients were then calculated by fitting the volume traces with a single-exponential curve and comparing the experimental time constants to time constants derived from one-parameter simulations.^37^ Therefore in our case, the difference in volume correlation between compounds in this proxy might lead to errors for the structurally diverse configurational isomers, especially that the permeabilities of nucleosides are usually considerably greater than something impermeable. Alternatively, several recent studies utilized fluorescent or isotope labeling to investigate the permeability of primitive molecules in vesicles.^40,54^ Nevertheless, the fluorescent labeling requires the modification of the original molecular structure, which might not reflect the actual permeability of a nucleoside or nucleotide in our study. The isotope labeling was monitored by scintillation counter, but again, the limitation is that only one penetrant can be labeled and tested at a time. A more operational and straightforward assay is desirable for evaluating the permeation of multi-component mixtures to identify the most penetrative isomer. Ideally, such an assay should offer high throughput and rapid quantification, such as liquid chromatography (LC), because our focus was on the permeability order rather than absolute values. The permeation of biorelevant molecules across biomimetic membranes may follow similar principles.^55^ Over the past two and half decades, the parallel artificial membrane permeability assay (PAMPA) has become a standard lipid-membrane-based platform, and gained increasing application of permeability measurements in drug discovery and formulation development.^56^ It is a well-established method for assessing the permeability of biomolecules across a membrane between donor and acceptor compartments (Fig. 2b). Given the difficulties of other methods for analyzing our mixtures, we found PAMPA to be a suitable alternative, as it is fast and capable of simultaneously handling mixed compounds in a single well.^57^ Therefore, we employed this assay to investigate the permeation order of nucleoside and nucleotide isomers.

To justify the utilization of PAMPA, we assessed its reliability by testing two reference solutes, with ribose being a representative non-ionic reference and 5-carboxyfluorescein being the ionic one. The results were compared with vesicle penetration data reported by Szostak,^37^ in which the same membrane component (oleic acid) was used. The permeability coefficient of ribose in PAMPA was 6.8×10^-8^ cm/s, slightly higher than in vesicle, but still within the same order of magnitude (Table 1). Similarly, the permeability of 5-carboxyfluorescein in PAMPA was 0.26% per day, closely aligning with the release rate of vesicle-encapsulated 5-carboxyfluorescein over the same time period (< 0.75% per day).^37^ Therefore, these data indicated that PAMPA offers a permeability level that is consistent with the vesicle method, qualifying it as a reliable alternative for studying the permeability of biomolecules.

**Table 1.**
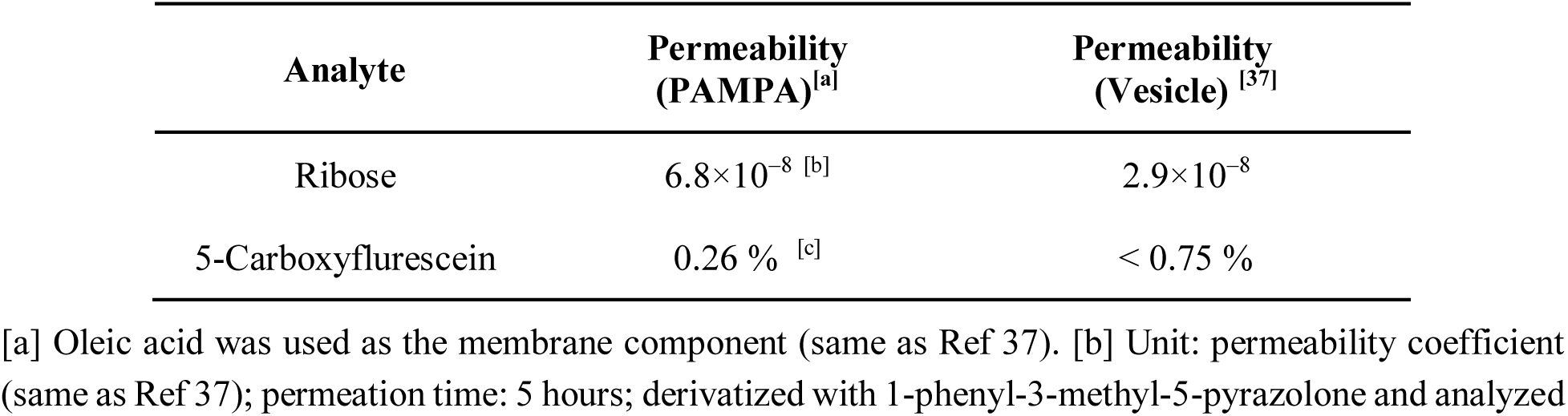
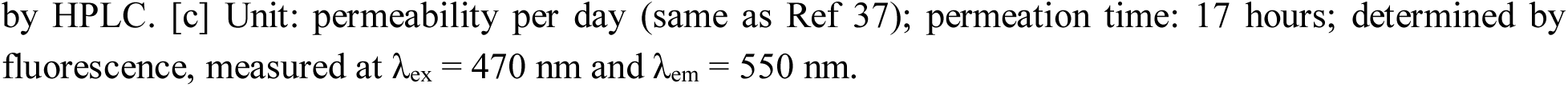
Comparison of permeabilities of ribose and 5-carboxyflurescein measured by the PAMPA and vesicle methods.

| Analyte | Permeability (PAMPA)<sup>[a]</sup> | Permeability (Vesicle)<sup>[37]</sup> |
| --- | --- | --- |
| Ribose | $6.8 \times 10^{-8}$ [b] | $2.9 \times 10^{-8}$ |
| 5-Carboxyfluorescein | 0.26 % [c] | $< 0.75$ % |

[a] Oleic acid was used as the membrane component (same as Ref 37). [b] Unit: permeability coefficient (same as Ref 37); permeation time: 5 hours; derivatized with 1-phenyl-3-methyl-5-pyrazolone and analyzed
by HPLC. [c] Unit: permeability per day (same as Ref 37); permeation time: 17 hours; determined by fluorescence, measured at $\lambda_{\text{ex}} = 470$ nm and $\lambda_{\text{em}} = 550$ nm.

The generation of a pH gradient across the lipid membrane is the foundation of energy transduction in modern cells.^58^ The importance of a transmembrane proton gradient has been addressed in developing both the early protomembrane models^59^ and the late ones.^60,61^ Szostak and coworkers pointed out that the growth of pure fatty acid membrane could generate a transmembrane pH gradient in the absence of metal cations, resulting in the acidification of the intracellular fluid.^59^ In contemporary cells, the cytosolic pH (∼7.2) is typically slightly lower than the extracellular one (∼7.3–7.4), and the pHs of many individual cell organelles are even lower. Therefore, we assumed that the extracellular fluid is near neutral (e.g. pH ≈ 7)^42^ while the intracellular fluid is weakly acidic (e.g. pH ≈ 6). To begin, the two mixtures of *N*9-ribosylated adenines and guanines (as **A_mix_** and **G_mix_**) were prepared via the FaPyA and FaPyG routes.^15,16^ For pyrimidine nucleosides, their limited availability hampered us from obtaining a complete set of the four *N*1-nucleosides. Instead, **C_mix_** and **U_mix_** were prepared by mixing the commercially available α/β*-*cytidines and α/β*-*uridines. To maintain consistency with the vesicle conditions, all solutes and solutions in this study were prepared in metal-free form. The permeation experiment was carried out in the polycarbonate (PC) 12-well microtiter filter plates at room temperature. The well was coated with lipid membrane (decanoic acid/decanol = 2:1 for **A_mix_** and **G_mix_**; or decanoic acid/decanol/glyceryl monocaprylate = 4:1:1 for **C_mix_** and **U_mix_**) and incubated for 20 min. Aliphatic alcohol or monoglyceride was frequently employed in reported studies as an essential co-simulant of protomembrane, since they help reduce the charge density of fatty acids,^62^ enhance permeability and the tolerance range of pH,^63,64^ cations^65^ and heat.^66^ The Szostak lab also reported that adding a nonionizable, alcohol/ester-based membrane component such as monomyristolein, can help stabilize the pH gradient across fatty acid membranes.^59^ Other membrane compositions might afford even greater permeability of the canonical nucleosides, but optimizing the lipid membrane for each compound was not the goal of this study. It was observed that excess amphiphiles (and alkane as loading solvent if needed) not retained in the pores immediately floated up upon contact with water. Consequently, only a minimal amount of amphiphiles is retained at the PC pores (see Supplementary Information for weight measurements) and self-assemble upon contact with water to form a bilayer,^56^ which has been verified by transmission electron micrograph (TEM). ^67^ We chose the minimum time required for the solutes in the acceptor compartment to become quantifiable by HPLC or extracted ion chromatography (EIC), when a clear baseline was achieved and peaks were integrable. This time was 48 h for purine nucleosides and 5–8 h for pyrimidine nucleosides. After that, the solution in the acceptor compartment was analyzed by HPLC or liquid chromatography-mass spectrometry (LC-MS), and carefully compared with the original mixture. Since the chromatograms for **C_mix_** and **U_mix_** were relatively clean, their compositions were determined by UV integrals. For **A_mix_** and **G_mix_**, due to the inseparable minor products from their syntheses, the EIC peak areas of [M+H]^+^ were used for quantifying their percent compositions, whose relative areas accord well with UV integrals.

The results for the permeation experiments are shown in Fig. 3–4. For **A_mix_**, **β*-*f-A** (β*-*adenosine, desired) gained 18% through permeation (almost doubled), the highest magnitude among the four isomers, and became the most abundant one in the acceptor plate. The order of percentage increase largely matched with the reverse order of RP-HPLC elution: **β*-*f-A** > **α*-*p-A** > **α*-*f-A** > **β*-*p-A** (Fig. 3). Notably, the once most abundant isomer (**β*-*p-A**, undesired) had the sharpest decrease by 20%. For **G_mix_**, the desired **β*-*f-G** (β*-*guanosine) also became the predominant species in the acceptor plate. Like **β*-*f-A**, the percentage of **β*-*f-G** also doubled through permeation, with the most drastic increase by 19%. Consequently, the percentage shares of the undesired guanosine isomers were brought down, with the largest drop by 19% for the formerly prevalent **β*-*p-G**.

**Fig. 3.**
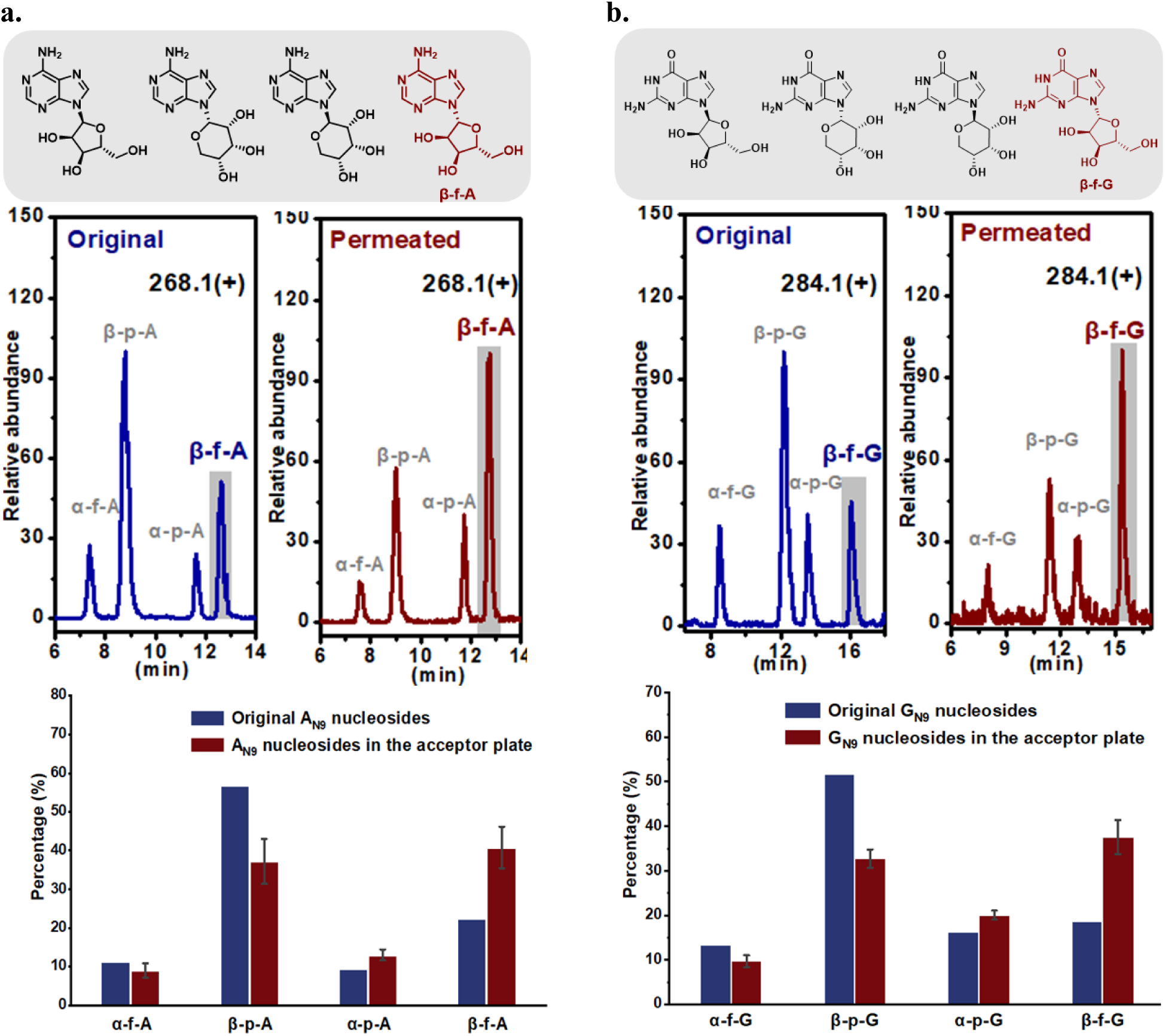
EIC chromatograms and compositions of the original purine nucleoside mixtures (blue) and the permeated ones (red): (**a**) adenosine isomers (**A_mix_**, searched for [M+H]^+^ = 268.1); (**b**) guanosine isomers (**G_mix_**, searched for [M+H]^+^ = 284.1). Membrane component: decanoic acid/decanol = 2/1. Permeation time: 48 h. Percentage compositions were determined by EIC integration. The percentage values of the acceptor compartments were based on three runs with error bars shown. The peaks of extant isomers are in the shaded region of the chromatogram.

**Fig. 4.**
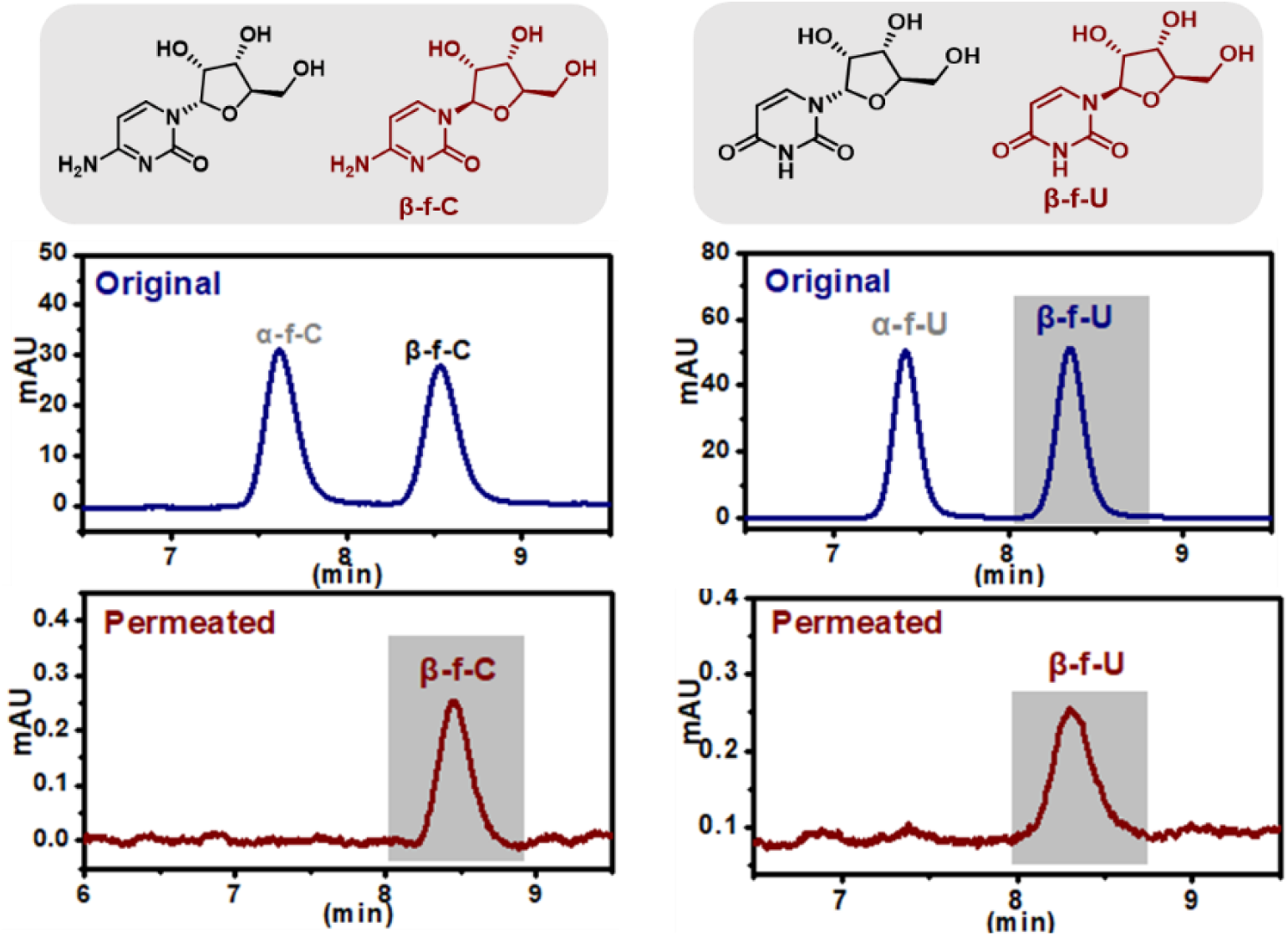
LC chromatograms of the original pyrimidine nucleoside mixtures (blue) and the permeated ones (red). (**a**) cytidine isomers (**C_mix_**); (**b**) uridine isomers (**U_mix_**). Membrane component: decanoic acid/decanol/glyceryl monocaprylate= 4/1/1. Permeation time: 8 h for **C_mix_** and 5 h for **U_mix_**. Percentage compositions were determined by LC integration. The peaks of extant isomers are in the shaded region of the chromatogram.

Furthermore, to evaluate the permeation behavior of **A_N9_** nucleosides in real cells, NCI-H1975 cells were incubated with the test solution (prepared by diluting the ribosylated-FaPyA mixture in PBS, pH 7.0) for 8 hours at 37 °C. After incubation, cells were thoroughly washed to remove extracellular residues, then mechanically disrupted to release intracellularly accumulated components. LC-MS analysis of the cell lysates revealed that the two pyranosyl isomers were selectively minimized through permeation, and **β-f-A** became the predominant isomer (see EIC traces in Figs. S32-1 and S32-2). This cellular experiment demonstrates that the selective permeation of β-furanoside is not limited to artificial lipid membranes but also operates in a genuine biological context, validating the physiological relevance of the proposed selection mechanism.

For **C_mix_** and **U_mix_**, because the initially attempted decanoic acid/decanol (2:1) membrane exhibited integrity issues with pyrimidine nucleosides (though no such leakage was observed in any other experiments), a more stable composition (decanoic acid/decanol/glyceryl monocaprylate (4:1:1)) was used instead. The extant nucleosides **β*-*f-C** (β*-*cytidine) and **β*-*f-U** (β*-*uridine) became almost the sole component through permeation (Fig. 4). Although calculations may not fully capture intrinsic properties of the analytes, we performed molecular dynamics (MD) simulation to rationalize the observed selectivity difference between pyrimidine and purine nucleosides. The free energy change (ΔG) for permeation of β-versus α-furanoside has been calculated (see Supplementary Information). The β-F anomer is more favorable by 0.12 kcal/mol in cytidines (**β*-*f-C**) and 0.28 kcal/mol in uridines (**β*-*f-U**). In contrast, **β-f-A** is only favored by approximately 0.02 kcal/mol. Although alternative membrane compositions might yield higher nucleoside permeability, the central aim of this study was to establish the order of permeation, rather than to pursue optimization of the membrane composition.

### Outward Permeation of Ribonucleotides: Selection of 5’-Nucleotide

In most examples of nucleoside phosphorylation, the resulting 5’-nucleotide is not exclusive, commonly with concomitant formation of 2’-and 3’-nucleotides.^13^ Given that the activation of 2’-or 3’-nucleotides leads to rapid cyclization,^68^ forming a 2’,3’-cyclic phosphate that traditionally cannot be incorporated efficiently during template copying, ^69^ it is possible that evolutionary selection favored 5’-nucleotide as oligomerization substrate for this reason. Although 2’-and 3’-nucleotide are ruled out for polymerization, they could be common products of RNA degradation. ^70^ Can these undesired isomers be transported out of a protocell? Again, simple biophysical factors could be responsible for that. With regard to the permeation of nucleotide, Szostak reported low permeability of 5’-AMP due to its charged and lipophobic nature, unless carrier cations such as Mg^2+^ are present which are disruptive to many protomembranes.^45^ Nevertheless, the permeability order of these nucleotide isomers still merits an investigation, which might give a clue to the understanding of the selection mechanism. Inspired by the elution order of nucleosides, we evaluated the retention behaviors of nucleotide isomers of all four genetic alphabets by reverse-phase HPLC. The results revealed a high regularity that 5’-phosphate (5’-nucleotide), with whichever nucleobase, had the *smallest* retention factor (Fig. 5). Evident differences of retention factor were observed among 5’-, 3’-, 2’-, and 2’,3’-cyclic phosphates. This result suggests that a separation mechanism, if ever existed, might preferably allow the permeation of the more lipophilic nucleotides. Therefore, it can be further inferred that 2’-, 3’-and 2’,3’-cyclic phosphates (if formed intracellularly, see further to the below) can probably be screened out by reversely crossing the protomembrane, whereas the desired 5’-phosphate tends to be trapped for longer within protocells.

**Fig. 5.**
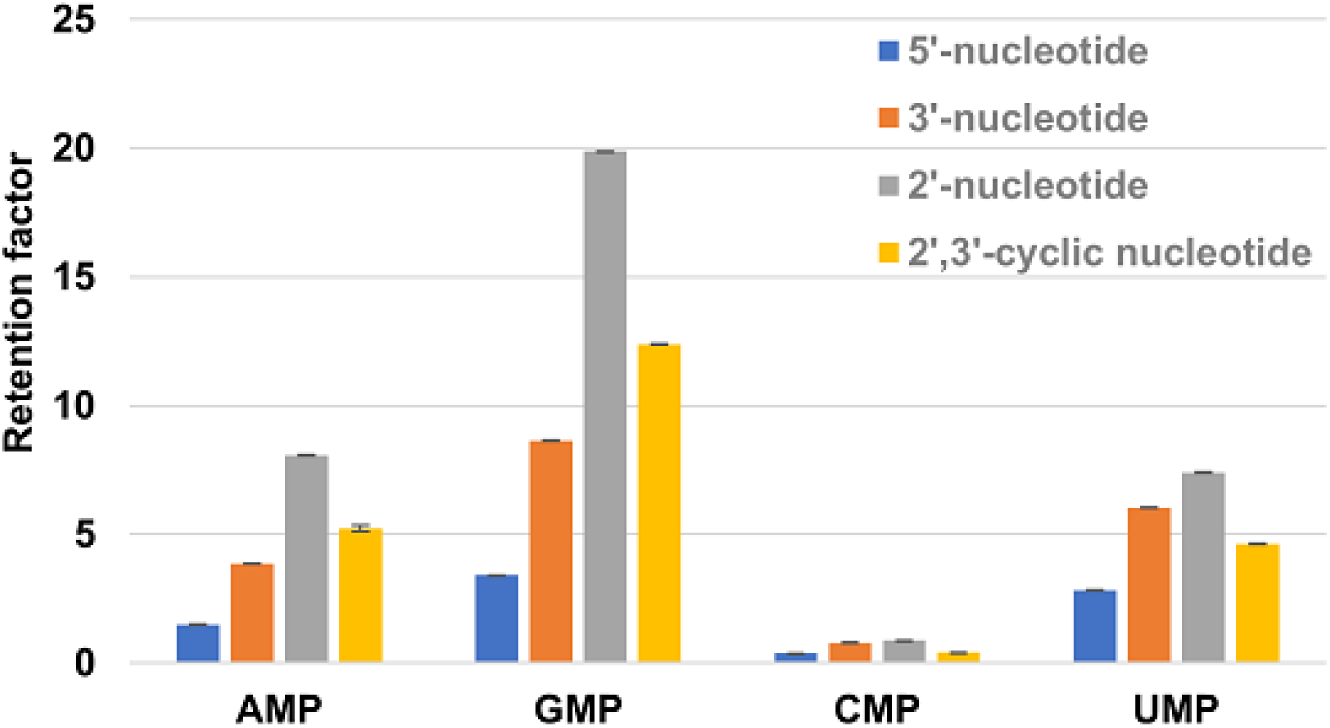
Retention factors of 5’-, 3’-, 2’-phosphates and 2’,3’-cyclic phosphate of all four genetic alphabets. Retention factors of nucleotide isomers were calculated by their retention times measured on a Shimadzu LC-2050 system with SymmetryShield^TM^ RP18 column (5 μm, 100 Å, 4.6 mm I.D. × 150 mm), at a detection wavelength of 254 nm, with water (adjusted with 0.1% formic acid) as the eluent at a flow rate of 1.0 mL/min. Measurements were based on three runs with error bars shown (with very tiny variation).

To test this assumption, an aqueous mixture of 5’-and 2’,3’-cyclic AMPs was subjected to the PAMPA assay with the PC 12-well microtiter filter plates (pH_donor_ = 6, pH_acceptor_ = 7). The well was coated with lipid membrane (decanoic acid/decanol = 2:1). The permeation experiment was run for 48 h. With analyses by LC, it was discovered that among the permeated nucleotides, the percentage content of 5’-AMP decreased by 27.4% as compared to the original solution, while 2’,3’-cAMP gained the same percentage (Fig. 6 and 7), which was consistent with the predicted order (2’,3’-cAMP > 5’-AMP). The results with GMPs, CMPs and UMPs followed the same pattern, with the 2’,3’-cyclic phosphate being more permeable. Owing to the ionic character of nucleotides, the extent of permeation was smaller than the corresponding nucleosides, thus peaks of trace precursory nucleosides and related impurities were amplified in the acceptor plate. It is worth emphasizing that although the examined conditions might not be comprehensive, we did not aim to seek an optimal lipid membrane for each group of nucleotides with higher permeability. It was also technically unrealistic to extend the permeation time to timescales relevant to geological ages, although it is reasonable to believe that longer periods would result in an increased number of permeated molecules. We also subjected the mixed monophosphates (5’-, 3’-and 2’-phosphate) for all genetic alphabets to the same assay. The permeability order was consistent with the retention factors. Noteworthily, 5’-phosphate was the only isomer with decreased percentage in the permeated samples for all four nucleobases (see Supplementary Information). In general, 2’,3’-cyclic phosphate experimentally demonstrates considerably greater permeability than all monophosphates, using 5’-phosphate as a reference.

**Fig. 6.**
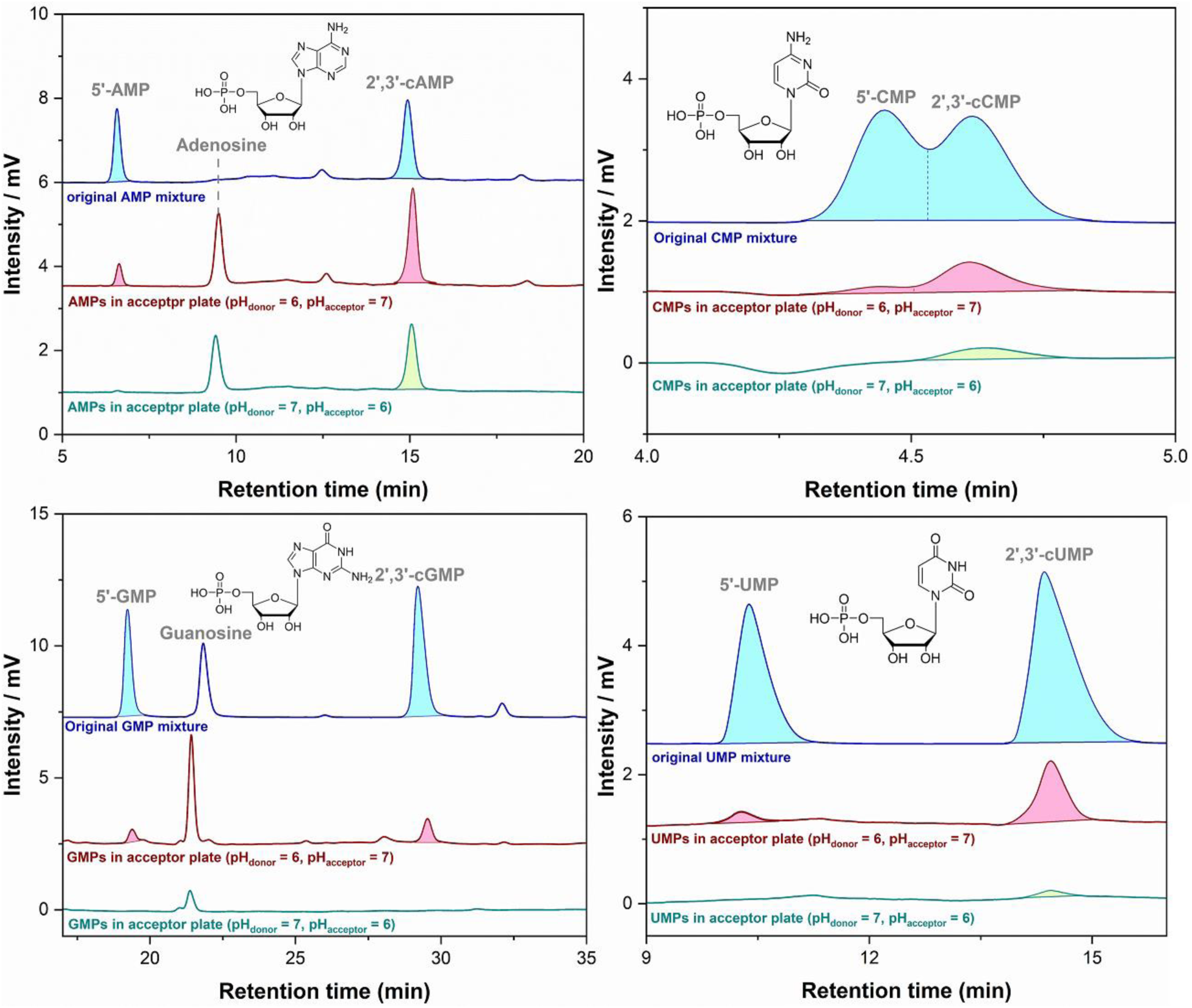
LC chromatograms of 5’-phosphate and 2’,3’-cylic phosphate in the original solutions (blue, upper) and the acceptor compartments (red, middle), and the results at the inverted pH values (green, lower). Membrane component: decanoic acid/decanol = 2/1. Permeation time: 48 h. Normal pH condition: pH_donor_ = 6, pH_acceptor_ = 7. Inverted pH condition: pH_donor_ = 7, pH_acceptor_ = 6. The peak of trace nucleoside or impurity in the mixture may be intensified in the acceptor plate due to higher permeability compared to nucleotide.

**Fig. 7.**
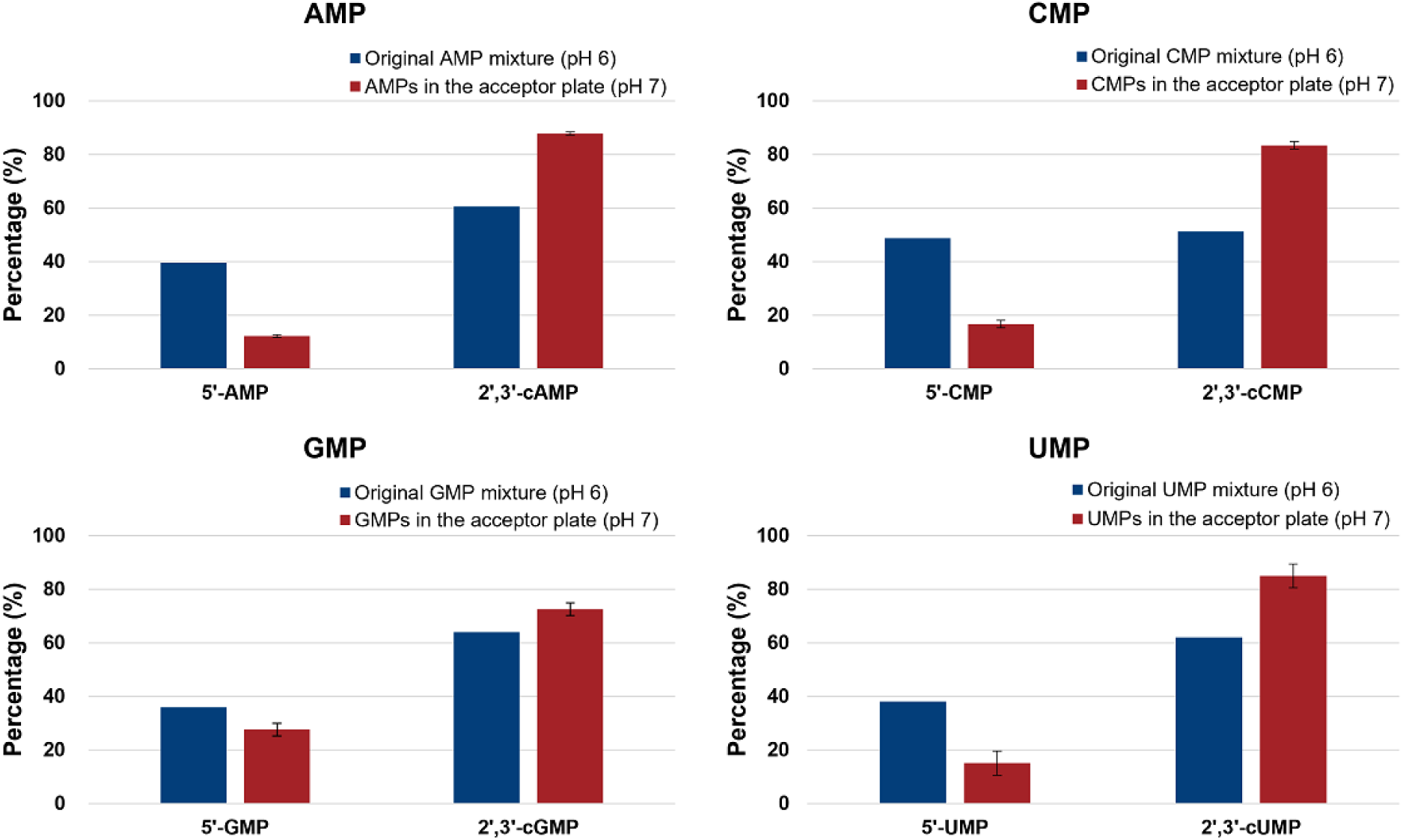
Change of isomeric compositions of 5’-phosphate and 2’,3’-cylic phosphate through permeation. Percentage compositions were determined by LC integration. The percentage values of the acceptor compartments were based on three runs with error bars shown.

Besides the primary observation, it was also essential to discuss other pH conditions. To that end, we assessed the permeabilities of nucleotides with inverted pH values of the donor and acceptor compartments (pH_donor_ = 7, pH_acceptor_ = 6). The results in Fig. 6 (bottom chromatogram in each group) evidently display the significantly diminished permeabilities of 5’-phosphate at the inverted pH, indicating that the influx of nucleotides is far more difficult than the efflux. These data are also supportive of a scenario that the phosphorylation of nucleoside possibly occurs intracellularly. Phosphoric acid ^71^ was proven permeable into the protocell at low concentration, and usable as a phosphorylation agent therein to react with the permeated β-furanoside (see Supplementary Information). We also examined systems with equal intra-and extracellular pH, representing scenarios where the pH gradient is absent or has been disrupted (e.g., by metal cations). The selective permeation of 2’,3’-cyclic phosphate was still observed (see Fig. S13).

Under these conditions, permeation remains effectively irreversible, as solutes that have entered the extracellular fluid are likely to diffuse away or be removed by flushing. Interestingly, at equal pH 7 on both sides, 5’-AMP became markedly less permeable than at pH 6, whereas 2’,3’-cAMP remained penetrative presumably because of its lower ionic character.

To rationalize the permeability difference among the nucleotide isomers, we first elected to evaluate the accessible non-polar surface area (ANSA) of the AMPs. Density functional theory (DFT) calculations were performed. The geometries of AMPs were optimized using the B3LYP density functional, with 6-31+G(d,p) as the basis set. Solvation effect in water was considered. The optimized geometries with electrostatic potential maps are displayed in Table 2. The ANSA values were calculated by subtracting the accessible total surface area (ATSA) with the accessible polar surface area (APSA). It turned out that 5’-AMP had the largest APSA (120.8 Å^2^) and the smallest ANSA (75.1 Å^2^). A possible underlying reason behind this observation is that 5’-nucleotide poses phosphate, the most polar moiety of the molecule, to be oriented outward (Table 2). In 2’-or 3’-nucleotide, the phosphate group is concavely positioned thus much less accessible than that of 5’-nucleotide, rendering them less hydrophilic. The ANSA results followed the order of 5’-AMP (75.1 Å^2^) < 2’,3’-cAMP (90.6 Å^2^) < 3’-AMP (94.2 Å^2^) < 2’-AMP (95.4 Å^2^). This calculated order showed better agreement with the retention factors than with the experimentally permeated amounts, because the calculated ANSA is based on an electronically neutral molecule (the net charge of a nucleotide at a given pH can be a non-integer, which cannot be assigned in the computation), and the retention factors are measured using acidic elution (isocratic water with 0.1% formic acid) which is near monophosphates’ isoelectric point. Experimentally, 2’,3’-cyclic phosphate is the most permeable. This observation is explained by the fact that at the near-neutral pH (6–7) of the permeation experiments, cyclic phosphate bears less negative charge than monophosphate. This result may also be reflected in the calculated APSA of 2’,3’-cAMP (109.1 Å^2^), which is the smallest of all AMP isomers (least polar).

**Table 2.**
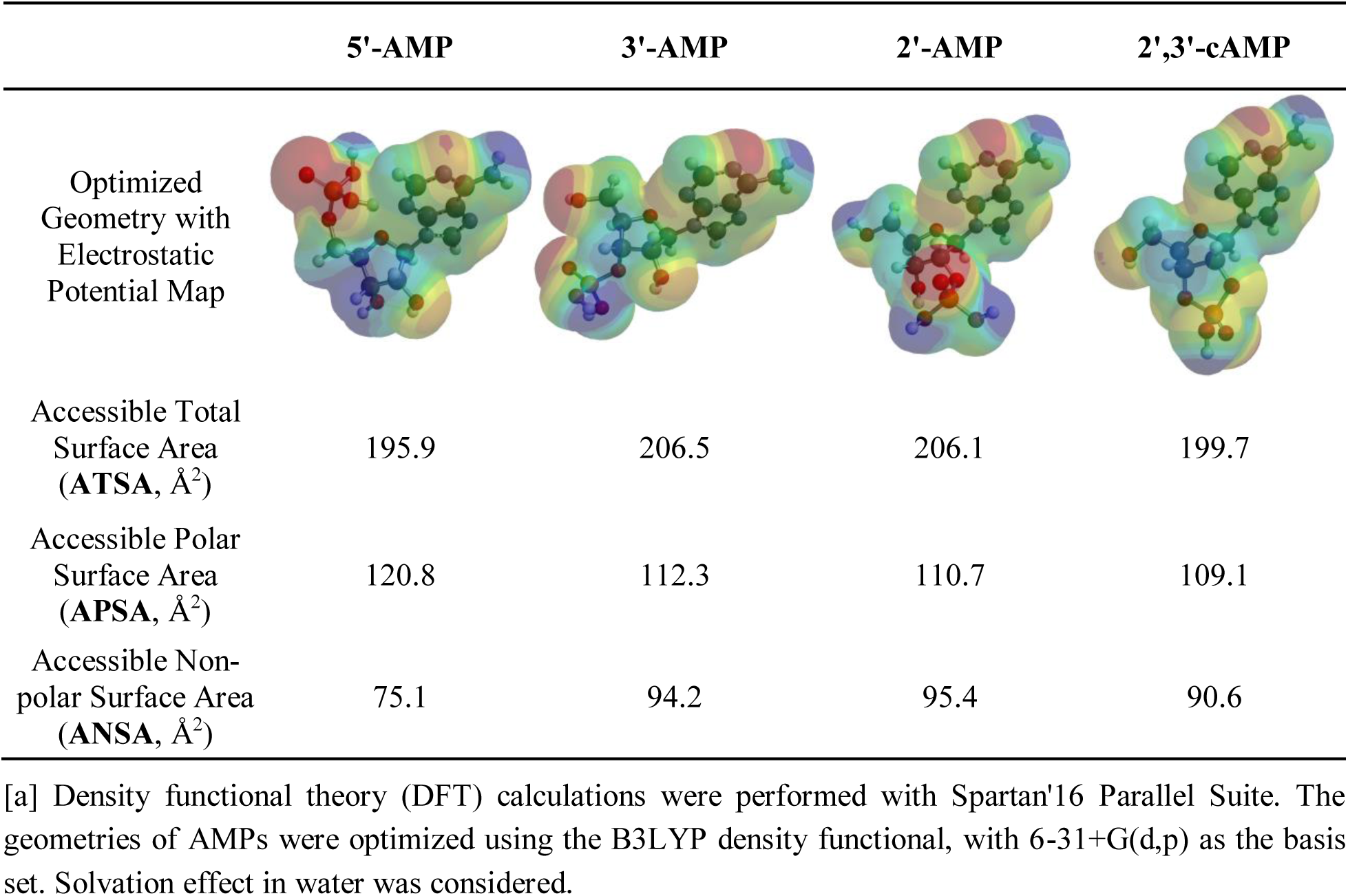
Optimized geometries of AMPs with electrostatic potential maps and the calculated surfaces areas.^[a]^.

|  | 5'-AMP | 3'-AMP | 2'-AMP | 2',3'-cAMP |
| --- | --- | --- | --- | --- |
| Optimized Geometry with Electrostatic Potential Map |  |  |  |  |
| Accessible Total Surface Area (ATSA, Å<sup>2</sup>) | 195.9 | 206.5 | 206.1 | 199.7 |
| Accessible Polar Surface Area (APSA, Å<sup>2</sup>) | 120.8 | 112.3 | 110.7 | 109.1 |
| Accessible Non-polar Surface Area (ANSA, Å<sup>2</sup>) | 75.1 | 94.2 | 95.4 | 90.6 |

[a] Density functional theory (DFT) calculations were performed with Spartan'16 Parallel Suite. The geometries of AMPs were optimized using the B3LYP density functional, with 6-31+G(d,p) as the basis set. Solvation effect in water was considered.

Apart from the DFT calculation, we were keen to explore the mechanism of nucleotides interacting with the lipid membrane in more detail, to see if there are any actual interactions taking place and why a selectivity is observed. Molecular dynamics (MD) is an effective technique for simulating molecular diffusion at the atomic level. Several previous studies have combined MD simulations with PAMPA experiments, ^72, 73^ merging diffusion analysis with free energy calculation. This integrated approach provides a comprehensive framework for investigating membrane permeability in greater detail. In this section, we used umbrella sampling simulations to calculate the free energy distribution of the potential of mean force (PMF) for the permeation of mono-and cyclic-phosphates (AMPs) across a lipid bilayer assembled by decanoic acid and decanol. The stabilization process was illustrated through a series of simulation snapshots (Fig. 8; see Supplementary Information for more details and data). Initially, decanoic acid, decanol and water were randomly arranged within the cubic box. Over time, they organized into a lipid bilayer, resembling a cell membrane. This result reinforces the reliability and plausibility of using decanoic acid and decanol as protomembrane components in the PAMPA experiments.

**Fig. 8.**
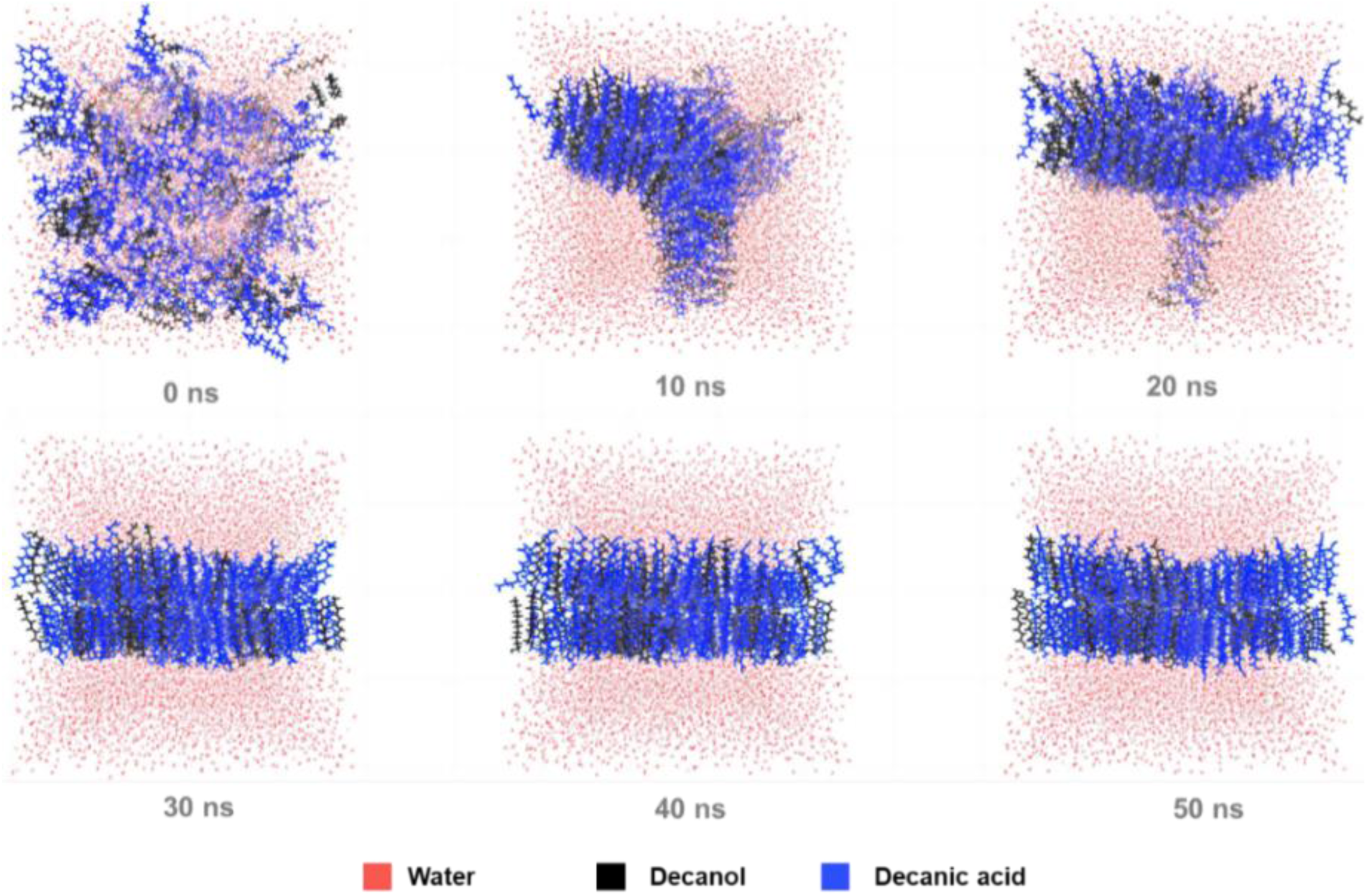
Assembly process of decanoic acid and decanol in the lipid-water system. MD simulations were carried out with GROMACS 2022.4, utilizing the GAFF force field and the TIP3P water model. The system contained 400 decanoic acid molecules and 200 decanol molecules, which were placed in a cubic box with the dimension of 8 nm × 8 nm × 8 nm.

The binding free energy changes of AMPs were reflected by the PMF curve. The calculated PMF curves for 5’-, 3’-, 2’-AMP and 2’,3’-cAMP across the assembled bilayer are shown in Fig. 9. The energy barriers for the AMPs were calculated as follows: ΔG_5’-AMP_ = +7.68 kcal/mol, ΔG_3’-AMP_ = +5.93 kcal/mol, ΔG_2’-AMP_ = +5.54 kcal/mol, and ΔG_2’,3’-cAMP_ = +6.53 kcal/mol. With the highest energy barrier, 5’-AMP exhibited the weakest permeability of all AMPs. The PMF results aligned well with the permeability order observed for the three monophosphates in PAMPA. Again, as the non-integer net charge of the nucleotides could not be simulated, the higher experimental permeability of 2’,3’-cAMP was not reflected in these results.

**Fig. 9.**
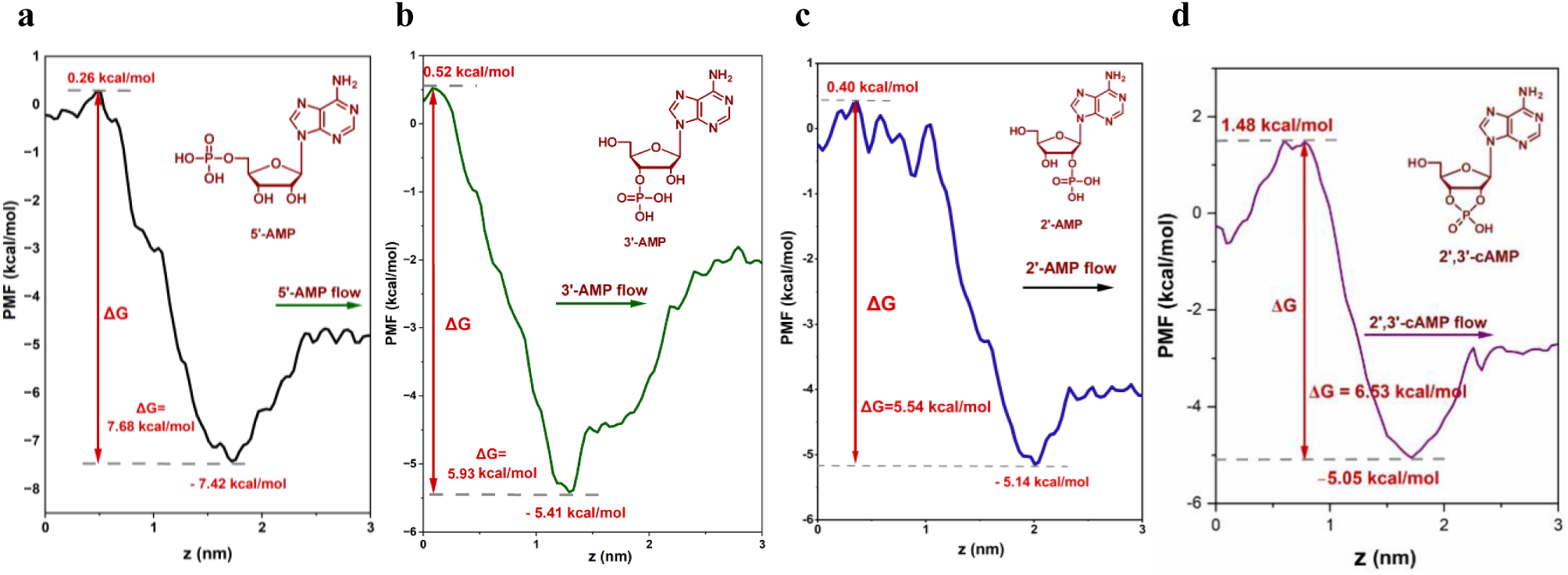
Calculated PMF curves for the mono and cyclic AMPs across the decanoic acid-decanol bilayer. (**a**) 5’-AMP; (**b**) 3’-AMP; (**c**) 2’-AMP; (**d**) 2’,3’-cAMP. Umbrella sampling was used to calculate the PMF free energy profiles for the three AMP molecules (see Supplementary Information for details). PMF profiles were then calculated using the weighted histogram analysis method (WHAM) implemented in GROMACS.

The experimental and computational results suggest that permeation is again a plausible mechanism for the selection of the desired configuration of RNA precursors, not only for nucleoside but also for nucleotide. Ideally, in the more basic extracellular fluid, the exited undesired nucleotides have little chance to reenter the protocell, rendering this process irreversible (Fig. 10a). Even without a proton gradient, the permeated undesired isomers outside the protocell are more susceptible to being flushed away by water currents than the retained 5’-nucleotide, making them unlikely to accumulate locally. Hence, the phosphorylation step can be advanced by the continuous removal of the undesired products out of the intracellular fluid. The unwanted nucleoside or nucleotide outside the protocell can be recycled back to their precursors via the hydrolysis of the glycosidic or ester bond. The intracellularly enriched 5’-nucleotide can be subsequently activated for nonenzymatic oligomerization.^74,75^ Although beyond the scope of this study, it should be noted that commonly proposed activating reagents for oligomerization such as imidazole,^76^ acetaldehyde and methyl isocyanide^74^ are relatively lipophilic, with calculated LogP ranging from –0.5 to around 0. In this study, the experimentally determined permeability data of canonical nucleosides and nucleotides are summarized in Fig. 10b. Apparently, the measured permeability (as LgP_app_, see Supplementary Information for the calculation) of an extant nucleoside (β-furanoside) is 0.5 to 1.0 order of magnitude greater than its nucleotide counterpart (β-furanoside-5’-phosphate), with the exception of β-guanosine which is over 10^2^ times more penetrative than 5’-GMP. Although it remains uncertain whether the tested lipid membranes resemble actual protomembranes on the primordial Earth, this observed trend suggests that nucleotides, unlike their nucleoside precursors, may tend to remain localized near their site of formation. This also implies that as polymerization proceeds, bulkier oligomers become progressively less likely to cross the protomembrane as their chains elongate.

**Fig. 10.**
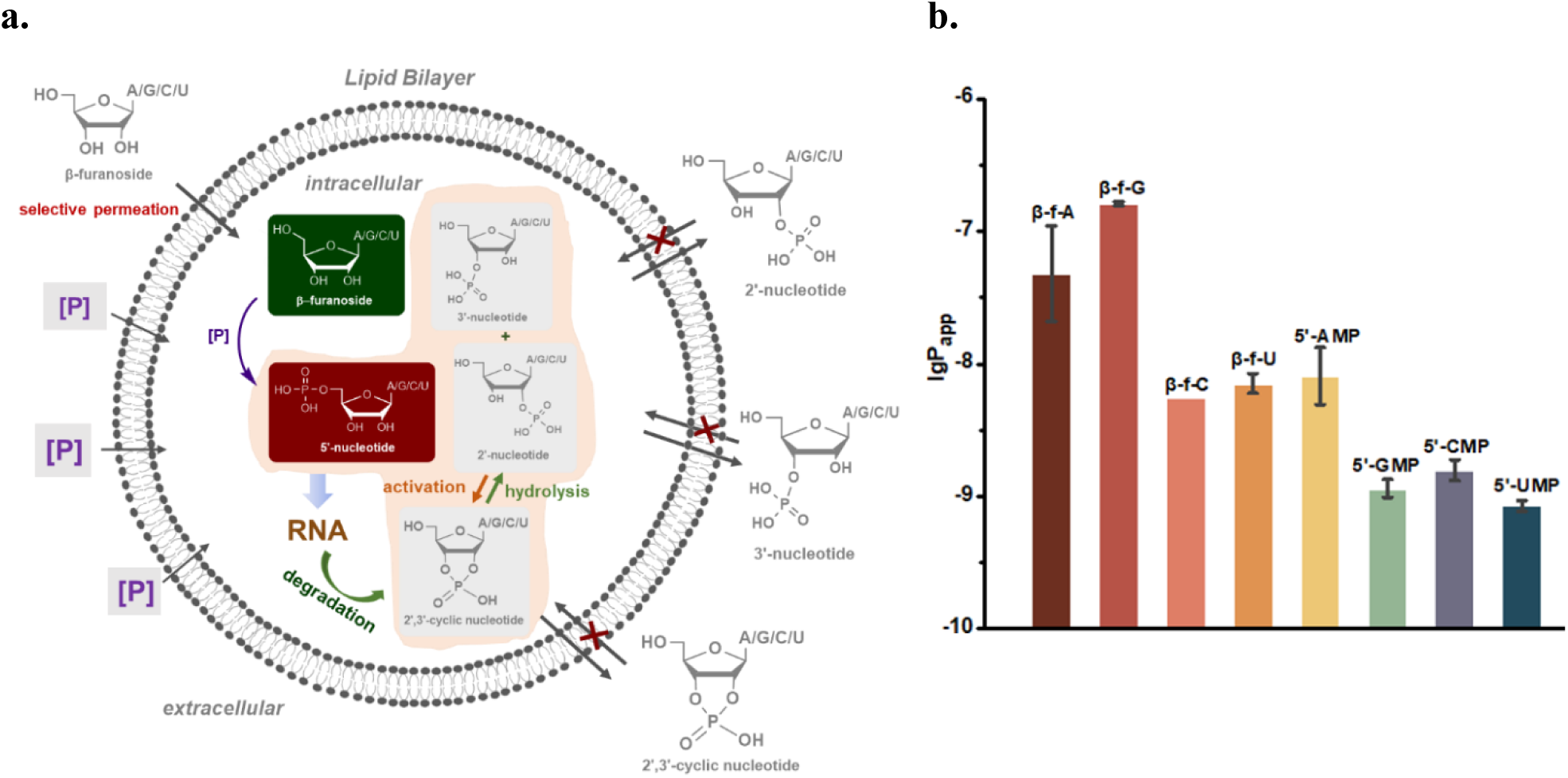
Selection of extant nucleoside and nucleotide configurations based on permeability. (**a**) Illustration of the membrane-facilitated selection of β-furanoside and 5’-nucleotide (“P”: phosphoric acid or phosphate); (**b**) Measured permeabilities of the extant nucleosides and nucleotides (measurements were based on three runs with error bars shown).

### Simulation of a Rapid Lipid-water Partition of Ribonucleosides with an Emulsifying Micromixer

Fatty acid and alcohol have been proposed as the main protomembrane components due to their structural simplicity and more dynamic behaviors relative to phospholipid. Their syntheses could be catalyzed by clay minerals.^49^ Subsequently, mineral surface adsorbs the formed fatty acids and provides nucleation sites for vesicle formation.^77^ Normally, the lipid bilayer of a vesicle is comprised of fatty acid and its conjugate base at a neutral to weakly basic pH. However, the primitive atmosphere rich of CO_2_ might constantly create an acidic environment on the early Earth,^78,79^ where the formation of the conjugate acid-base pair became difficult if not sufficiently buffered. Thus, this scenario challenged us to consider what would happen when lipids in acidic form interact with nucleosides/nucleotides in a non-vesicle, oil droplet state. ^80^ Analogously, examples of substances stored in oil droplet are commonly found in modern biology. In eukaryotic organisms, lipid droplets serve as essential organelles for storing triglycerides and cholesterol esters to produce membrane building block.^81^ It can be inferred that under a primordial scenario, water containing nucleosides/nucleotides frequently flushed over the mineral surface, mixing the solutes with the amphiphiles. The coarse surface of riverbed or floodplain might have enhanced the mixing as a static mixer.

To this end, we specifically designed a micromixer aided by computational fluid dynamics (CFD), to gain an insight into the transport of solute to lipid phase. The core of the micromixer comprises arrayed microstructured units to mimic the uneven mineral surface (Fig. 11). The aqueous solution of nucleosides and the mixed lipid (decanoic acid: decanol = 2:1) were introduced into the micromixer via two pumps (see Supplementary Information). The lipid-water biphasic flow was cleaved by the shear force at the sharp microstructure of the first unit, and continued to be split into smaller droplets in the repetitive units (see the Supplementary Video file). With an inner volume of only 1.73 mL, the micromixer was able to afford a thoroughly mixed fluid in only 0.94 second of residence time. The linear flow rate was approximately 1.6 m/s, making it a proper simulant for a natural stream. Interestingly, it was clearly observable that the mixture was emulsion-like near the mixer inlet, and became much more transparent at the exit, indicating the formation of smaller oil particles.

**Fig. 11.**
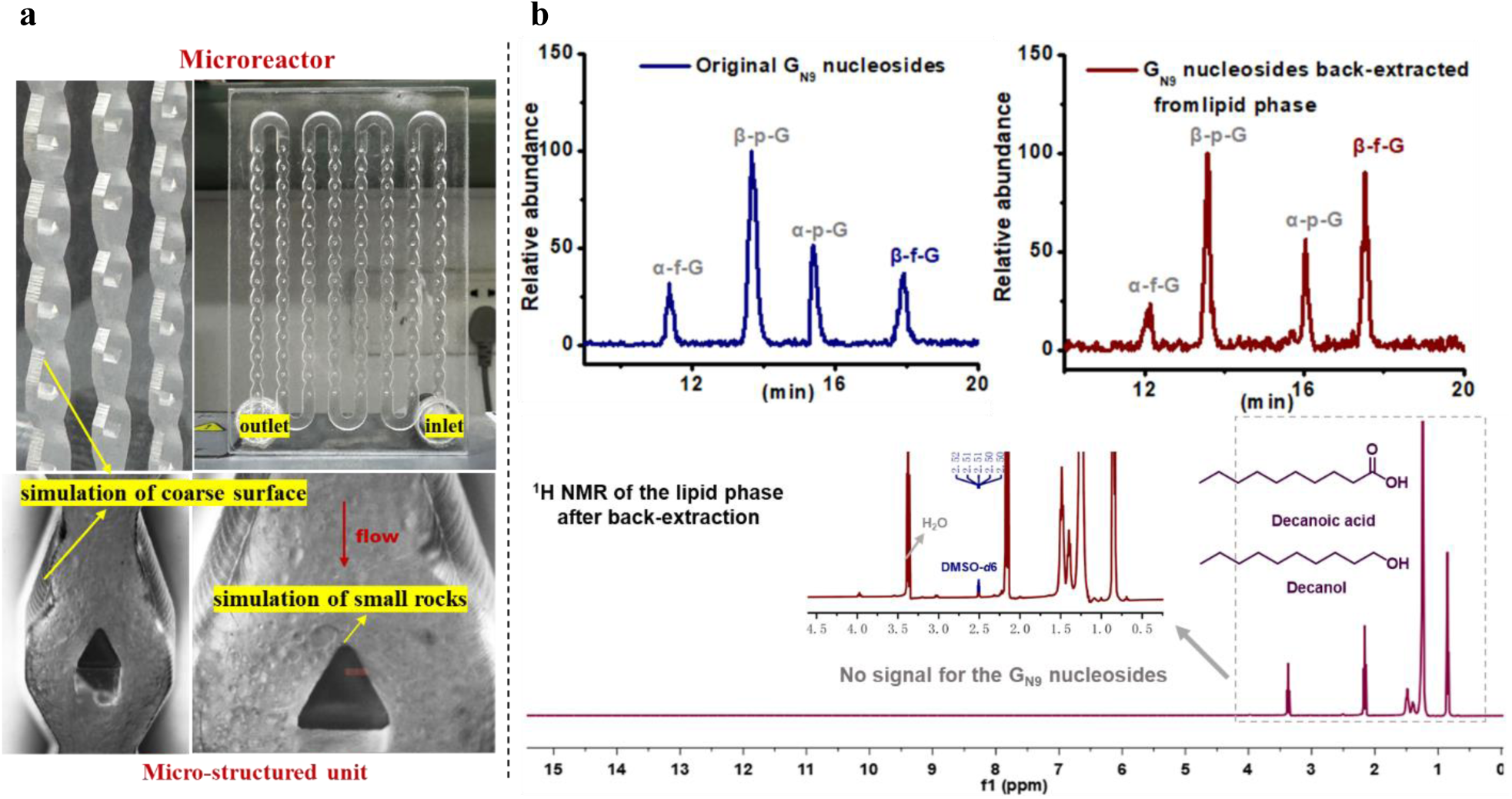
Simulation of a rapid lipid-water partition of nucleoside with an emulsifying micromixer. (**a**) Mixing the aqueous solution of **G_N9_** nucleosides (100 mL/min) and lipid (decanoic acid/decanol = 2/1, 10 mL/min) in a micromixer with arrayed sharp microstructures to mimic an uneven mineral surface (see the Supplementary Video). The micromixer was made by polycarbonate (PC) using laser engraving. (**b**) EIC of the original and back-extracted (from lipid) guanosine isomers and the ^1^H NMR of the lipid phase after back-extraction (frequency: 400 MHz; solvent: DMSO-*d*6, 2.50 ppm; the water signal (3.33 ppm) was embedded within the triplet at 3.37 ppm).

The collected fluid was centrifuged and separated. To explore what might occur when lipid droplets retaining nucleosides collide with incoming fresh water, the separated lipid phase was back-extracted with water, and the latter aqueous solution was instantly analyzed by LC-MS. For **G_mix_**, the change of isomeric distribution was close to the permeation experiment — the percentage of **β-f-G** rose most significantly through lipid partition (Fig. 11). Similar results can be obtained in batch, where the aqueous solution were vigorously stirred with lipid for 14 hours, followed by a back-extraction. These results validated a scenario that β-furanoside can be at least temporarily preserved and enriched in oil droplets, which is likely followed by continuous water flushing and extraction. ^1^H NMR of the remaining lipid phase after back-extraction showed no signals related to either the guanine ring or the ribose skeleton, which excluded the possibility of the nucleosides getting stuck in the lipid (Fig. 11). Furthermore, it also verified that the same lipid components in the PAMPA experiments did not retain any penetrant when exposed to water. In summary, β-furanoside can be selected, no matter the amphiphile assembles into lipid membrane or droplet. It could selectively traverse the water-lipid interface and enrich over time, preparing to engage with the subsequent transformations towards RNA.

## CONCLUSION

In conclusion, we have developed a rational model that promotes the selection of β-furanoside and 5’-nucleotide, the required configurations for the core structure of extant RNA. The selection mechanism is based on permeability differences between desired and undesired isomers across a lipid barrier. Permeabilities of nucleoside and nucleotide isomers were extensively investigated. β-Furanoside is more capable of crossing a lipid membrane, whereas nucleosides with other configurations are less permeable. Nucleotides undergo a reverse screening at the lipid membrane, which selectively allows undesired isomers to pass through. Through these inward and outward permeation processes, β-furanoside and β-furanoside-5’-phosphate become enriched. This study supports a plausible scenario in which selection of the canonical RNA structure can be achieved by a post-synthesis separation process, which helps eliminate undesired isomers, yielding purer materials for subsequent synthesis and replication.

Multiple selective factors, such as stability, synthesis and separation, may have collectively led to the selection of the desired precursors for extant RNA. However, additional insights can be gleaned from the data in this study. One such insight is that these observations likely reveal a diversification-selection evolutionary pattern at the molecular level. In particular, the results suggest that a synthetic route need not be stringently selective. Contrary to some traditional beliefs, we propose that a competent primitive synthetic process might simply generate a broad set of products to achieve sufficient diversity, rather than affording the desired product in high yield and selectivity. In a much less demanding yet still effective manner, selection likely occurred after a synthetic process rather than during it. These findings also provide evidence of protocellular substance exchange in the primordial world, driven purely by biophysical forces to advance critical steps in the progression of primitive biomolecules, which could be one of the earliest protocellular functions.

## ACKNOWLEDGMENTS

This work was supported by the National Natural Science Foundation of China (21872068, X.W.).

## DATA AND MATERIALS AVAILABILITY

The authors declare that all data supporting the findings of this study are available within the paper and its supplementary information files. The Supplementary Information available for this Article contains all procedures, NMR spectra, HPLC traces and LC-MS data, computational details, and a Supplementary Video file.

## AUTHOR INFORMATION

Correspondence and requests for materials should be addressed to Xiao Wang.

## AUTHOR CONTRIBUTIONS

X.W. conceived the project. Z.-R.Z., Q.-Q.C. and B.-Y.Z. performed the experiments, analyses and characterizations. H.-X.X. and Q.-Q.C. analysed the data. C.-C.G. ran simulations and calculations. X.W., Z.-R.Z. and Q.-Q.C. wrote the paper with suggestions from all authors.

